# Investigating motor preparatory processes and conscious volition using machine learning

**DOI:** 10.1101/2020.09.07.286351

**Authors:** Siobhan Hall, Dawie van den Heever, Mikkel C. Vinding, Linzette Morris

## Abstract

**Background:** Conscious volition is a broad term and is difficult to reduce to a single empirical paradigm. It encompasses many areas of cognition, including decision-making and empirical studies can be done on these components. This work follows on the seminal work of Libet *et al*. (1983) which focused on brain activity preceding motor activity and conscious awareness of the intention to move. Previous results have subsequently faced criticism, particularly methods used to average out EEG data over all the trials and the readiness potential not being present on an individual trial basis. This following study aims to address these criticisms.

**Objectives:** To use machine learning to investigate brain activity preceding left/right hand movements with relation to conscious intent and motor action.

**Methodology:** The data collection involved the recreation of the Libet experiment, with electroencephalography (EEG) data being collected. An addition made in this study was the choice between “left” and “right” while observing the Libet clock to subjectively mark the moment of conscious awareness. Twenty-one participants were included (four females, all right-handed). A deep (machine) learning model known as a convolutional neural network (CNN) was used for the EEG data analysis.

**Results:** Subjectively reported conscious intent preceded the action by 108 ms. The CNN model was able to predict the decision “left” or “right” as early as 4.45 seconds before the action with a test accuracy of 98%.

**Conclusion:** This study has shown motor preparatory processes start up to 4.45 seconds before conscious awareness of a decision to move.

## 1. Introduction

The question around conscious volition has been topical for millennia, especially considering its links to moral responsibility and the ownership of that responsibility. This question ultimately seeks to give, or take away the agency (or ownership) to the self and to our decision-making processes. However, volition is a broad term, and cannot be easily reduced to a single experimental paradigm. There are different definitions to account for (and little consensus), not to mention different aspects of cognition to accommodate (Roskies, 2010). Roskies (2010) proposes that there are five aspects to be considered when developing a more complete understanding of volition. These are: *intention, initiation, decision-making, feeling and executive control*. These are not distinct properties, but rather overlapping. However, separate investigations can contribute to the overall understanding. A free choice, or a free decision making process requires options, to avoid being denounced as the result of ‘unspecific neural preparatory action’ in the brain (Soon *et al*., 2008). There is the specific requirement of being able to do otherwise, should one choose to (Dias & Lavazza, 2016). The alternative to this is a completely random, or irrational decision, which would typically present as a reaction or urge, and not an act of volition. Further, these free choices also need to be carried out in the real world, in real world situations, with real world consequences, but, also most importantly the decision needs to be bereft of any external or internal processes that the person themselves is not in direct control over (Maoz *et al*., 2019). The latter is not always possible in a controlled experimental setting, but this is the requisite to which empirical studies should scale.

Predominantly a debate nested solely in philosophy, one of the first empirical investigation into the question of volition was in 1983 by a team of researchers headed by Benjamin Libet. Following on from the work of Kornhuber and Deecke, who discovered the so called “bereitschaftpotential” in 1965, otherwise known as the “readiness potential” (RP); Libet designed an experiment to study this sub-conscious neural precursor (Kornhuber & Deecke, 1965). This “readiness potential” is a neural precursor to movement and is found in the averaging of electroencephalography (EEG) data over many trials. EEG is the recording of the brain’s electrical potential and forms the basis of data collection in the Libet study as well as this study. The RP presents itself in the EEG data before movement. The seminal work of Libet *et al*. set out to determine whether or not this readiness potential occurs before or after the person is consciously aware of their intention to act. (Libet *et al*., 1983)

The Libet paradigms consists of the following set up. Participants were asked to use the “Libet clock” to report the moment that they became aware of their “intention” or “urge” to move. The moment of conscious awareness of intention has been termed “W” and the awareness of movement relative to the movement itself, i.e. the “M” moment. The results of this study found the RP to be present 550 ms before actual movement (“M”), while the onset of the conscious awareness of the intention to act (“W”) was found to occur 200 ms before movement (“M”). This means, there is a 350 ms period between the RP, the neural indicator for preparation of movement, and the moment of conscious awareness (“W”) – i.e. 350 ms period of brain activity without our conscious awareness, or involvement. (Libet et al., 1983).

The trivial separation of the subconscious (or, as referred to in this study, the pre-conscious) and the conscious is not as easy as assumed by Libet *et al*. (1983) with the use of “W” to delineate the moment of pre-conscious intention entering conscious awareness of the intent. This is a poor reduction of the complex phenomenon that is volition. There have been subsequent studies, looking to build and improve on the foundation of Libet (Fried *et al*., 2011; Soon *et al*., 2008, 2013), addressing criticisms and contributing to the understanding of volition. These criticisms are still prevalent, specifically the readiness potential, the single finger movement and the subjective nature of reporting the moment of conscious awareness (Banks & Isham, 2008; Fried *et al*., 2011; Lau *et al*., 2004; Miller *et al*., 2011; Murakami *et al*., 2014; Schurger *et al*., 2012; Soon *et al*., 2008, 2013; Trevena & Miller, 2010).

The rise in neural activity before conscious awareness (i.e. the readiness potential [RP]) was prematurely taken as proof the pre-conscious is ultimately in control of our actions (Libet, 1999). The timing of the RP was also considered insignificant as it occurred hundreds of milliseconds before the action as well as being the result of averaging out over many trials. This averaging “collapse[s] the dynamic information” in the EEG – essentially taking out the unique differences in different trials and uniqueness in individual participants. This limits the potential to make inferences about the brain processing as we have only kept that which is common (average) among all trials (Delorme & Makerig, 2004). Further criticism was that the experiment involved an abrupt finger flexion and/ or wrist movement (Libet et al., 1983) – and this cannot be considered anything more than an urge. A free choice requires options, as well as consequence to be considered anything more than an automatic reaction (Soon *et al*., 2008). There is the specific requirement of being able to do otherwise, or choose another course of action, in the event the exact same situation could be replicated (Dias & Lavazza, 2016). It is for this reason choice will be included in the Libet paradigm. Trevena & Miller (2010) sought further insight into the readiness potential, suggesting that perhaps the RP was not specific to movement preparation, but rather it is an indicator of “on-going [background] brain processes” where attention specifically is employed. Therefore, the RP is not specifically “event-related”, but a rather vague signal and the claims of Libet et al. (1983) cannot hold true. Murakami *e al*. (2014), Schurger, (2018) and Schurger et al. (2012) proposed a model to explain the shape of the RP in that the result of random fluctuations crossing a threshold in order for an action to occur (stochastic accumulation model). It must be noted that the recent findings of Travers, *et al*. (2020) has shown the RP is not a product of random neural fluctuation crossing a threshold in order to be carried through, as initially proposed by the aforementioned authors, however, it still holds true that the RP is not a source of relevant information when investigating neural processes before voluntary actions. Miller *et al*. (2011) found the RP to be a mere artefact of monitoring the Libet clock. An artefact in EEG is a signal that arises for reasons other than consistent cerebral activity - such as a physiological heartbeat or blink, or in this case a non-physiological extraneous source: the monitoring of the clock used in all Libet paradigm-styled experiments (Schomer & Lopes da Silva, 2011). A final, but important criticism, is the recording of the moment of conscious awareness (“W”). This retrospective subjective report has been proven to be inaccurate and vulnerable to manipulation (Banks & Isham, 2008; Lau *et al*., 2007). Banks & Isham (2008) found that an audio or visual stimulus within 80ms of “W” can change the perception of time, to the degree that the participant could report “W” to occur *after* the action. This shift in perception could be as much as 44ms. The inaccuracy in the report is also likely due to the fact an agent (participant) does not have access to the temporal location of the “W” moment (Bayne, 2011). The retrospective and subjective nature of “W” will not be addressed further in this study; however, one can be more certain that the pre-conscious brain activity precedes conscious intent if there is a larger time period between these two occurrences. Explicitly studied reaction times (the difference between thought and action) by Thomson *et al*., (1992) found reaction times to be between 140 and 200 ms depending on the stimulus. Finding features in brain activity relating to decision-making significantly earlier than 200 ms before conscious awareness of intent can help compensate for the inaccuracies in marking the exact moment of conscious awareness.

In 2008, Soon *et al*., sought to determine the areas of the brain responsible for determining movement on a pre-conscious level, and how early this process begins. Instead of the traditional Libet paradigm, the researchers had the participants watch a stream of letters in order to mark the moment of awareness of intent to click a button with either their left or right index fingers (“W”). This addition of a choice brings us closer to a truly volitional action (Dias & Lavazza, 2016). Using functional magnetic resonance imaging (fMRI), the researchers were able to decode information about decision making in the pre-frontal and parietal cortices. The authors were able to predict which finger would move between seven and ten seconds before it entered the participants’ conscious awareness using multivariate linear classification. However, it must be noted this classification had an accuracy of 60 %; which is only slightly better than chance. This study was followed up in 2013, by Soon *et al*., in that instead of an arbitrary “left” or “right” decision, participants were asked to choose between two basic arithmetic operations – addition or subtraction. Besides aiming to see if prediction was possible, they were also looking for neural overlap (or lack thereof) between pre-conscious signals for decisions for movement and decisions for abstract intentions. The analysis of results focused on two determining points – the prediction of the timing of the decision as well as the prediction of addition or subtraction (“when versus what”) decisions. The build-up of “when” neural activity was found in the medial frontopolar cortex as well as the posterior cingulate cortex (precuneus specifically). The build-up of “what” neural activity was in the pre-SMA, the same area as the predictive neural activity for voluntary movement. In both cases, accurate predictions could be made as to whether the participant would perform addition or subtraction (“what”) and *before* (“when”) the participants themselves were consciously aware of this decision (Brass & Haggard, 2008).

In 2011, Fried *et al*. used a more invasive technique (depth electrodes) to study neurons during “self-initiated movements”. The study focused on neuronal activity in the SMA, pre-SMA and the AAC as well as 259 neurons in the temporal region. The participants followed the Libet paradigm, the only difference being the participants were asked to press a key when they felt the urge to do so. “Progressive neuronal activity” was observed 1500 ms before the participant’s report of making the decision to move, and the researchers could predict the movement 700 ms before this moment of decision (“W”). They concluded monitoring of only 256 neurons in the supplementary motor area is necessary to make this prediction.

The focus of the current study includes elements of the Libet paradigm, in order to address the criticism of Libet *et al*. (1983) as well as certain limitations of Soon *et al*. (2008, 2013): fMRI has a poor temporal resolution due to the low temporal resolution of BOLD (Blood oxygenation level dependent) signals. Further limitations, going beyond the Libet paradigm, include the fact that the majority of research using EEG is centred around acquiring methodology from previous investigations. In doing so, research loses innovation and becomes stagnant in terms of simply “confirming” the results of previous studies (Johannesen *et al*., 2016). This limits the analysis process – EEG has *millisecond* temporal resolution – ensuing innumerable time-points and features to be accounted for – and these purely neuroscientific methods end up excluding massive portions of these dynamic, cognitive components during analysis (Johannesen *et al*., 2016). Further, simple visual analysis of EEG data (as an applicable example), by a human observer introduces observer bias. Bias is introduced with the standard EEG analysis, viz. the averaging of the data across many trials. The averaging smooths out the signals to reveal ERPs (event-related potentials) – essentially taking out the unique differences in different trials (Delorme & Makerig, 2004). In other words, the RP (an example of an ERP) is not present on a single trial basis and averaging loses valuable cognitive information present in individual trials. Tenable conclusions cannot be drawn without all the information present in the EEG. Perhaps something we see as noise, because of prior human knowledge, may be a key neural marker in understanding something unprecedented about the brain.

It is for these reasons that machine learning is proposed as a tool to address the limitation of the RP. Deep learning is a branch of machine learning. The difference is the use of artificial “neural networks” (not to be confused with neural networks in the brain) which create depth in the model. “Deep” neural networks (DNN) are essentially a series of interconnected steps between input and output. This creates a “deep” network – with stacked algorithms between input and output. A convolutional neural network (CNN) is therefore used in this study.

Motivations for using a machine learning (ML) system include their ability to handle “fluctuating environments” (Géron, 2017), a key requirement in the context of EEG data analysis. It cannot be guaranteed that everyone’s EEG patterns will be the same, nor can it be assumed that each trial for one participant will be identical, due to confounding factors of trial-to-trial variations such (of varying origin) in the EEG data. ML algorithms have also shown their potential to gain insights into complex problems especially those problems in which there is “no good solution at all using a traditional approach” (Géron, 2017), giving us an avenue for innovative research and discovery.

This potential for innovation is essential in the context of this study. Neuroscientific research looking into the timing of conscious awareness of the intention to act has maintained a consistent approach since the seminal work of Libet *et al*. (1983). Machine learning is useful in its ability to aid pattern recognition, an attribute useful when the goal is to extract features in rich datasets such as EEG. This is in contrast to the traditional approach of investigating *a priori* evoked signals. The RP limits the analysis of the EEG to a single signal; while ML is preferable as it opens the analysis to include all the EEG without collapsing any dynamic information in the brain (Delorme & Makerig, 2004).

The current study therefore aims to address some of the criticisms and limitations of previous studies by adding a choice to the Libet paradigm and using machine learning for the analysis of the EEG data prior to conscious awareness of the decision to move either left or right. This study focuses on the motor preparatory processes relating to decision-making and the conscious awareness of volition

## 2. Method

The aim of this study is to accurately classify a decision (left or right) by employing supervised learning neural networks with EEG data from before an action and conscious awareness of a decision to move as the input.

### 2.1. Design

Data were collected from 21 volunteer participants (four females, all students) recruited from Stellenbosch University, South Africa. One participant’s EEG data were rejected on account of excessive EEG artefacts present within the data. All participants were right-handed. The age range was 18 to 28 years, with a mean age of 21.1 years (standard deviation ± 2.69). Each participant was screened according to the inclusion and exclusion criteria for their eligibility to participate in the study. The basis for inclusion was normal, or corrected-to-normal vision. Grounds for exclusion were predominantly based on medication, recreational drug use and/or prior head traumas, as these have been shown to alter EEG signals (Banoczi, 2005; Broglio, *et al*., 2011; Powers, *et al*., 2014; Schomer & Lopes da Silva, 2011; Teel, *et al*., 2014; Yang, *et al*., 2015). Informed consent was obtained in English from each participant prior to the trial. There was no financial compensation. The study received ethical and institutional clearance from Stellenbosch University (Health Research Ethics Committee: N18/04/048); having adhered to the ethical guidelines and principles of the international Declaration of Helsinki, South African Guidelines for Good Clinical Practice and the Medical Research Council (MRC) Ethical Guidelines for Research.

### 2.2. Data collection

The EEG data were recorded using the BRAIN Products ActiCAP EEG system with 128 gel electrodes. The Cz channel was the online reference and an average reference was used offline (Teplan, 2002). The data were collected at a sampling rate of 500 Hz and then downsampled to 256 Hz. A modified Libet clock (Vinding *et al*, 2013) was presented using the PsychoPy2 (Peirce *et al*., 2019; Peirce, 2007) coder functionality onto the computer screen. Each “second” on the clock represents 42.7 ms in reality (Libet *et al*., 1983). Participants were seated 60 cm from the 21” LCD monitor with a 1024×768 resolution and 60 Hz refresh rate.

All participants were instructed to follow a procedure similar to the Libet paradigm, including the subjective report of the moment of conscious awareness (“W”), with the main diversions from the original paradigm as follows: Participants were asked to click a button with either their left or right index fingers at will. A choice is important to not have a single action denounced as an unspecific urge or reaction. This methodology is similar to that of Soon et al. (2008) with the exception of EEG data being collected instead of fMRI.

### 2.3. Data pre-processing

All 128 channels of the EEG data were pre-processed. Pre-processing was performed for two types of analysis: (1) an event-related potential (ERP) analysis was done in order to recreate the RP as found in the original Libet paradigm as well as (2) the data was prepared as input to the convolutional neural network (CNN) used in the deep learning analysis. EEG pre-processing was performed using the open source software, EEGLAB (Delorme & Makeig, 2004) in the MATLAB environment (2019a, The MathWorks, Inc.).

The automation of artefact removal needs to be reconsidered. The use of automatic algorithm leaves little room for intuition regarding changes to the data. This is especially important to consider as these algorithms were designed with specific assumptions in mind – i.e. for more ‘pure’ neuroscientific analysis such as the averaging for ERPs; and not for machine learning. Machine learning requires an inherently different approach. It is for these reasons that the “manual” pipeline as implemented by An, *et al*. (2019) and Anders *et al*. (2018) was followed to account for these differences in assumptions and approach. One author was responsible for implementing the entire pre-processing pipeline for all participants

There are four main steps in removing noise from the data. These are listed as follows:

a. *Filtering*: to ensure only the frequencies of cognitive interest are included (i.e. high pass filter applied at 1.5 Hz and low pass filter at 50 Hz, i.e. frequencies from delta to gamma frequency waves).
b. *Bad channel rejection*: excessively noisy channels can skew all other channels measured relative to it as the chosen referencing technique is average referencing
c. *Artefact removal*: this is the process of removing non-stereotypical, non-recurring artefacts
d. *Independent component analysis (ICA)*: this is the process of removing stereotypical, recurring artefacts (non-cognitive components) such as eye blinks, the heart rate (pulse) and some muscular activity.

### 2.4. CNN analysis

The aim of this study is to determine how early motor preparatory processes start in our decision-making process. A high test classification accuracy using only EEG data from before conscious awareness of a decision to move would suggest there are features in the pre-conscious data relating to the decision, indicating a large role of the pre-conscious in our decision making.

The entire EEG montage of 128 channels were input to the CNN. The EEG data were segmented into windows. These EEG data were segmented into incrementally longer windows to observe the changes in accuracy over the different time stamps. However, only a fraction of these windows was fed into the algorithm, in order to determine the earliest time range required to make accurate classifications of the data.

Figure 1 illustrates the segmentation of data more clearly. It is important to note that only the time axis was segmented, not the channels.

**Figure 1:**
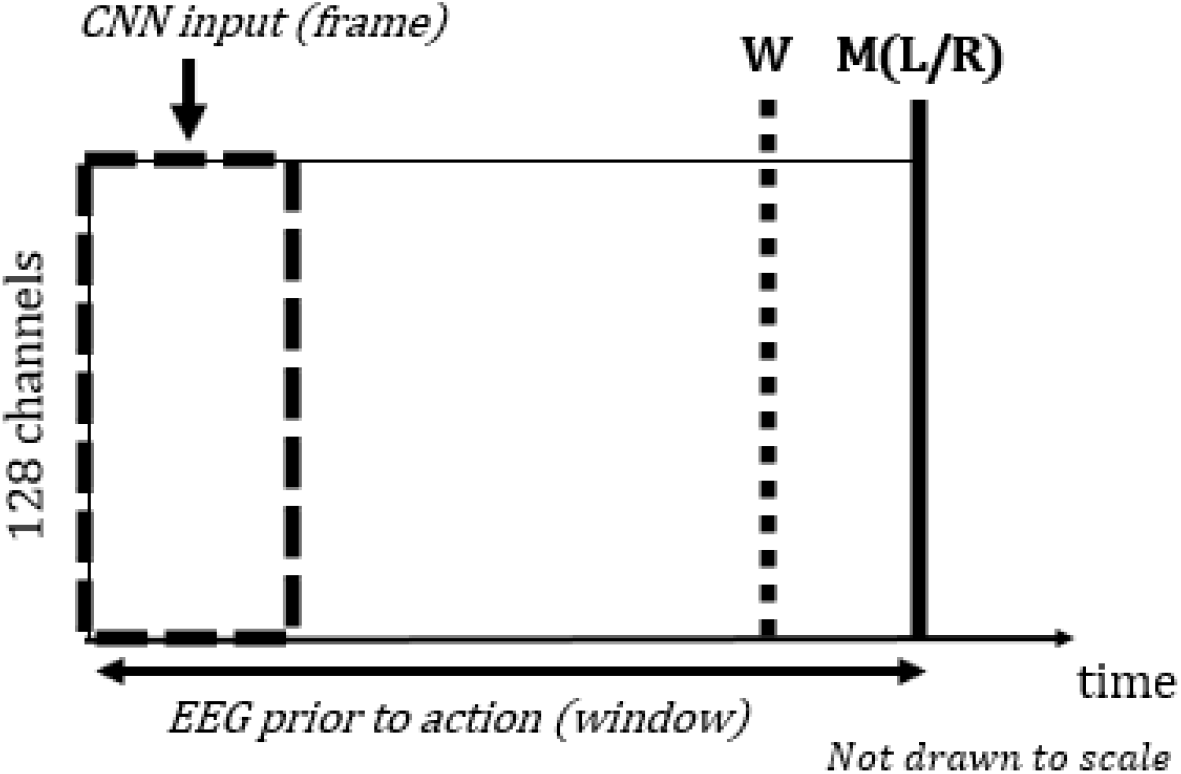
A depiction of the segmentation of the data into windows (EEG data prior to the action, labelled “M”) and subsequent frames as input into the CNN. “W” refers to the moment of conscious awareness

Table 1 summarises the timestamps of EEG data fed into the CNN model. The window refers to the period of time before the action (“M”), while a frame refers to the duration of the window fed fed as input into the CNN. Window durations were chosen to obtain highest classification accuracies as early as possible, while mitigating overlapping of the time frames. These frames ensure there is only data prior to conscious awareness of the intention to move.

**Table 1:** The timestamps of the data input to the CNN and respective number of samples

| Duration of the windows<br>(ms) | Duration of frame as CNN<br>input (ms) | Time stamps of the frame<br>(ms) | Total number of trial<br>samples available |
| --- | --- | --- | --- |
| 1500 | 375 | 1500:1125 | 2318 |
| 3500 | 1000 | 3500:2500 | 1962 |
| 4500 | 1250 | 4500: 3250 | 1791 |
| 5000 | 550 | 5000:4450 | 1680 |
| 7000 | 1875 | 7000:5125 | 1460 |

A trial sample, hereinafter referred to as a trial, is defined as one free willed decision, i.e. the start of the Libet clock to action that is recorded. Only trials that were longer than 2000 ms were included. In each respective duration, there were variable numbers of samples (albeit evenly distributed between “left” and “right” decisions) included for training, validation and testing. The variability in the numbers of these samples was due to the variability in the length of each trial, and some trials were shorter than others as it was purely dependent on the participants’ volition and the respective timing thereof.

Once each window was segmented into the relevant frame, the data was randomly split into the training, validation and tests sets. This random split ensures data from each participant were present in each respective set. The split is 70 % training data and 15 % for the validation and test sets, respectively. The test set is kept aside until after tuning of the hyperparameters in the CNN using the validation accuracies. This is because the purpose of the test data is to evaluate the deep learning model’s ability to perform on unseen data.

The final stage, having split the data, is to check the distributions of each label (i.e. “left” an “right”) in each subset. The model assumes a normal (consistent) distribution across all the sets (train, validation and test sets). Should the distribution be imbalanced (number of trials in one class outweighing the other) additional steps would need to be considered such as weighting the losses or down sampling the larger class by randomly deleting trials. The model used in this study was a convolutional neural network (CNN). The architecture was based on that of Schirrmeister, *et al*. (2017), developed specifically for EEG data, by adapting the state of the art AlexNET CNN used for image recognition. The model is coded using the high level neural network platform, Keras (Chollet, 2018). The basic architecture is that of a four-layer convolutional neural network, with one dense layer. The ReLU activation unit was chosen to be used in the hidden layers, with softmax used in the output layer. A dropout probability value of 50 % was used (i.e. each unit had 50 % chance of being included in that relevant iteration). Hyperparameter adjustments were made to the model using only the validation accuracy. The model is only deployed on the test set once for each window size. The test set is fed into the CNN model only once – as the final measure of the model’s performance. The test subset is akin to data the model will encounter when released into production (for example) and the test set results shouldn’t inform any hyperparameter tuning, as this can result in the model simply overfitting to the test set as well.

### 2.5. ERP statistical analysis

Further analysis was performed on the readiness potential to provide comparative measures between machine learning and the conventional neuroscientific methods. The aim of these ERP statistical analyses are to determine whether or not the the RP can be used to determine, or predict an action before it takes place.

#### 2.5.1. Determining the time of the RP onset

The RP investigated in this study is based upon a decision to move. Therefore, the RP onset was further analysed on an inter-subject basis to investigate the null hypothesis that there no significant difference between the “left” and “right” RPs at the Cz channel. There are different proposed methods to determine the RP onset (Verbaarschot *et al*., 2015). Two methods were employed in this research:

a. *Visual method:* The data were visually inspected, stepping from the moment of the action (time = 0ms) in the negative direction. Three consecutive time points where the EEG data above the baseline are averaged for the RP, per participant. (Keller & Heckhausen, 1990; Libet et al., 1983; Verbaarschot et al., 2015)
b. *T-test:* The datapoints were compared to the baseline using a t-test, starting from time = 0 ms and moving in the negative direction. Three consecutive datapoints with a p-value <0.05 were taken as the RP onset. (Verbaarschot et al., 2015)

#### 2.5.2. Non-parametric cluster based permutation tests

Non-parametric cluster based permutation tests (Maris & Oostenveld, 2007). The tests were conducted in two parts:

a. *Single channel data:* The RPs for each decision (“left” / “right”) were analysed on an inter-subject basis analysed to investigate the null hypothesis that there no significant difference between the “left” and “right” RPs at the Cz channel (critical alpha < 0.05). The input data were −1500 ms – 0 ms.
b. *Entire EEG montage across all windows used in the ML analysis:* The cluster based permutation tests were performed on an inter-subject basis across the entire montage (128 EEG channels by time) to test the null hypothesis that there no significant difference between the “left” and “right” RPs at each electrode site (critical alpha < 0.05) in the same windows used for input to the CNN (see Table 1). The tests were repeated for the five windows of EEG data that were used for input to the CNN. These durations correspond with the durations in the first column of Table 1. Cluster analysis improves the sensitivity of the test to changes in the spatiotemporal domain of the RP.

## 3. Results

### 3.1. Results from the convolutional neural network

The model was able to consistently classify the action “left” or “right” with accuracies of 99% using variable frames of data between 1.1 and 4.5s before the action took place. The results for each respective frame are presented in Table 2.

**Table 2:** Results of the mean test accuracies.

| Time stamps of data input into CNN (ms) | Test accuracy (%) of the CNN model | No. training samples | No. validation (and test) samples |
| --- | --- | --- | --- |
| 1500:1125 | 99.43 | 1622 | 348 (348) |
| 3500:2500 | 98.98 | 1374 | 294 (294) |
| 4500: 3250 | 99.63 | 1253 | 269 (269) |
| 5000:4450 | 98.6 | 1176 | 252 (252) |
| 7000:5125 | 50.45 | 1022 | 219 (219) |

Figure 2 shows the CNN model’s test classification accuracy at 1125 ms before the action. This illustrates the temporal position of this accuracy relative to the onset of the RPs (for the left/right decisions) and the “W” time. The RP onset (t-test) corresponds to the results in Table 4.

**Figure 2:**
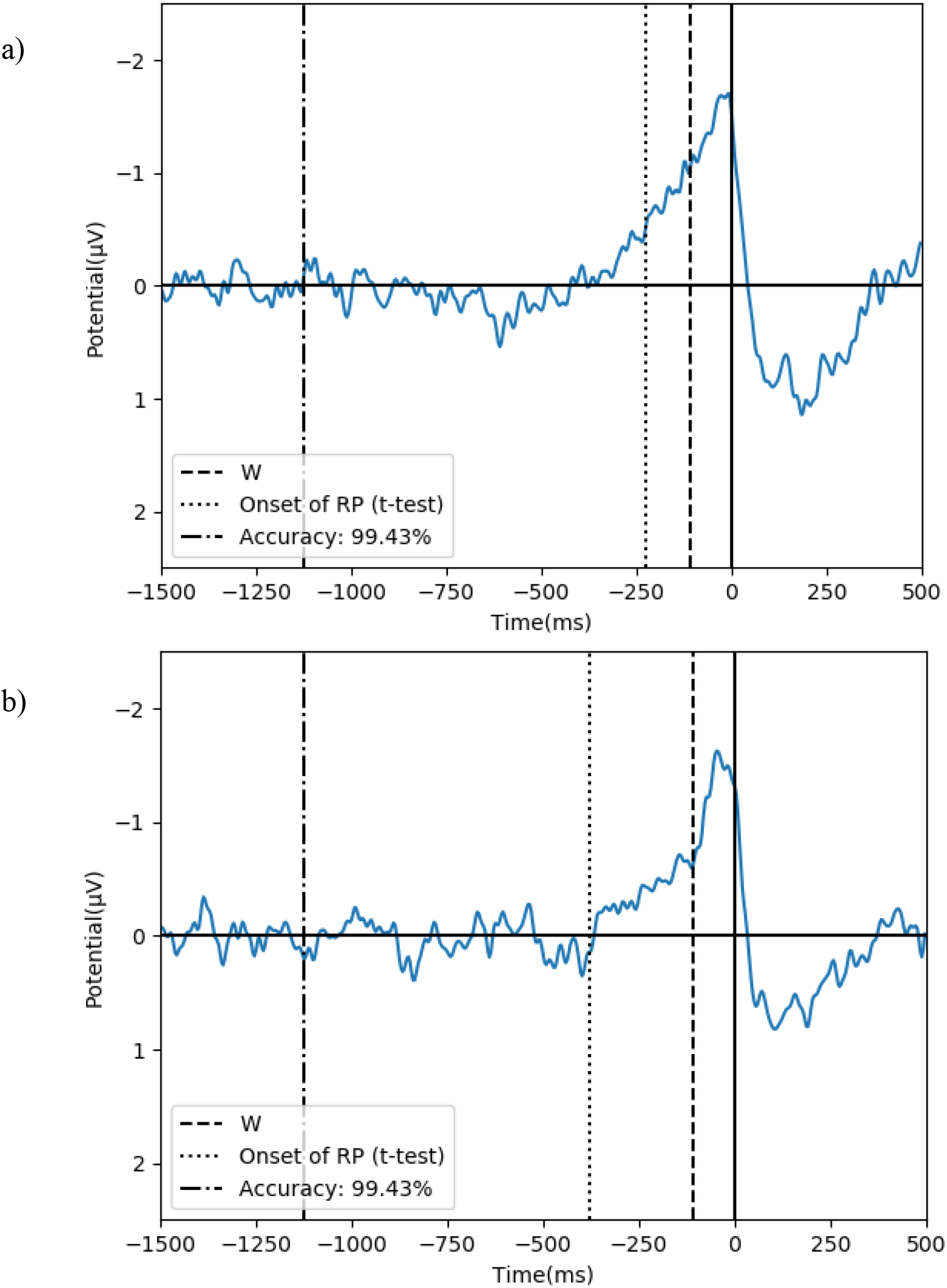
The comparison of the machine learning classification accuracy to the onset of the RP for the left decision (a) and the right decision (b)

### 3.2. Readiness potential to prove data validity

The first result is proving data validity. The graphs in Figure 3 present the averaged RPs for the “left” decision (a) and the “right” decision (b). No smoothing has been applied and these are from the Cz channel. The zero line indicated the moment of movement (“M”) and the “W” time is marked by the dashed line.

**Figure 3:**
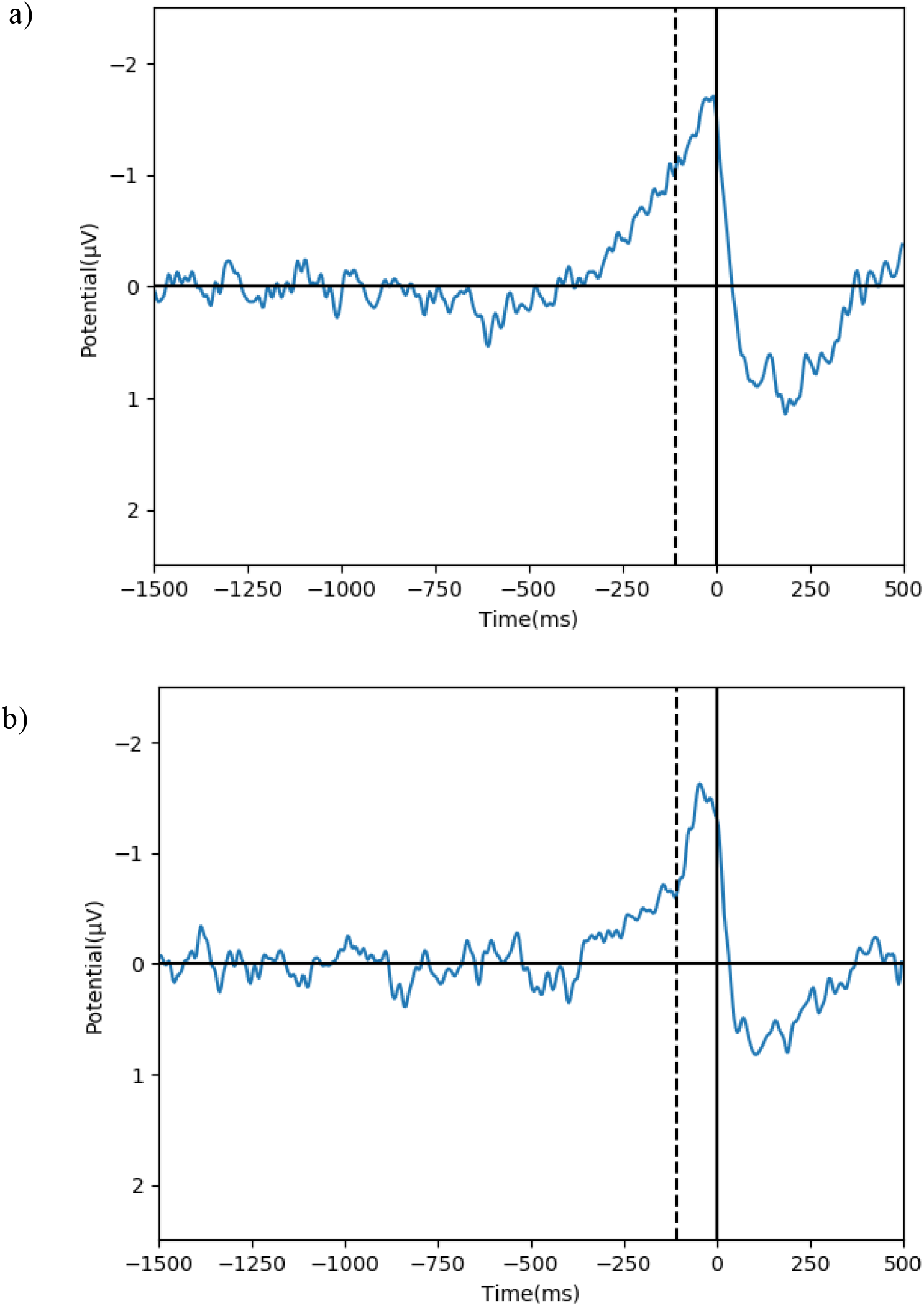
The reproduction of the readiness potential for the “left decision (a) and the “right decision” (b)

In this study, the subjective report for the moment of conscious awareness (“W”) was found to be an average of 108 ms before the action (standard deviation ± 0.11). This average is over all trials, regardless of the decision regarding the movement. In both RPs we see the rise in negative potential around 400 ms before the action, and 300 ms before “W”.

### 3.3. Comparison of the left and right readiness potentials

The following presents the results of the statistical analyses of the “left” and “right” readiness potentials.

#### 3.3.1. Determining the onset of the RP

a. *Visual test:* Results as presented in Table 3.
b. *T-test:* Results as presented in Table 4.

**Table 3:** Results of the visual test to determine average onset time of the RP.

| Action | Average time (ms) before action (0 ms) | Standard deviation |
| --- | --- | --- |
| Left | 214 | $\pm 126$ |
| Right | 180 | $\pm 113$ |

**Table 4:** Results of the t-test to determine average onset time of the RP at the Cz channel.

| Action | Average time (ms) before action (0 ms) | Standard deviation |
| --- | --- | --- |
| Left | 226 | $\pm 218$ |
| Right | 378 | $\pm 494$ |

#### 3.3.2. Non-parametric cluster based permutation tests

a. *Single channel data:* A significant difference between the “left” and “right” RP was found just before the movement (−70 to −200 ms). This finding is consistent with the lateralised readiness potential typical of the RP.
b. *Entire EEG montage across all windows used in the ML analysis:* The results are presented are Table 5.

**Table 5:** Results of the cluster-based permutation tests for the window timestamps as input to the EEG

| Time stamps of data input to the cluster-based analysis | Cluster p-value | Reject/ accept $H_0$ |
| --- | --- | --- |
| 1500:1125 | 0.004 | Reject |
| 3500:2500 | 1.000 | Accept |
| 4500: 3250 | 0.006 | Reject |
| 5000:4450 | 0.749 | Accept |
| 7000:5125 | 0.356 | Accept |

## 4. Discussion

Developing a firm understanding of the movement preparatory processes and conscious volition links to an understanding of moral responsibility and who, or what, should be assigned agency for our actions. In 1983, Benjamin Libet *et al*. introduced the Libet paradigm as an attempt to bring empirical evidence to support the debate about free will which previously centered around theoretical philosophy (Libet et al., 1983). Although the rise in neural activity before conscious awareness (i.e. the readiness potential) was taken as proof the pre-conscious is ultimately in control these conclusions were premature and a poor reduction of volition. This result has faced much criticism, and subsequent studies have attempted to address this.

The first step in analysing the data was confirming that the data collected in this study was representative of the data from the original Libet experiment (Libet et al., 1983). Obtaining this result was necessary to prove the validity of the experiment conducted in this research. Successful reproduction of the readiness potential (RP) at the Cz electrode for both the “left” and “right” decision was achieved. In confirming the validity of the EEG data, further comparison can be made in terms of addressing the criticisms around the Libet paradigm.

Having confirmed the representativeness of the data, we showed that it is possible to predict the decision to move the left or right hand from EEG data using a convolutional neural network, up to 4.34 seconds before the subjective reported time of the intention “W” and 4.45 seconds before the action. This shows there are neural preparatory processes ∼4 seconds before the conscious awareness of a decision to act, giving rise to evidence that the perception of conscious volition appears *after* the intention to act. Further, this action involved a choice between “left” and “right”. This study suggests the notion that there is neural information highly predictive of a free choice present before the subjective report of conscious awareness of a decision to move by participants. Of equal importance, and in accordance with the critique of classical event-related potential (ERP) methods and use of RP in these experiments, this study shows a prediction of the decision can be made long before the onset of the RP.

This is further confirmed by the results of the RP onset analysis tests. The visual test showed the RP onset to be, on average, around 200 ms before the action. The t-test yielded more variance in terms of the RP onset times, with standard deviations greater than ± 200 and a difference of more than 100 ms between the onset times for “left” and “right” RPs. The large standard deviation and inconsistent times for the RP onset in the respective decisions indicates large variability between participants, and these are not precise methods to be used to predict the RP onset relative to the action.

The RP is the result of averaging over trials and over participants. The RP is not present on an individual trial basis and not sensitive to unique dynamics of individual participants. As an average, it is not representative of brain dynamics at the points of consideration and it cannot be proven that the RP is in fact present *before* conscious awareness of volition on an individual trial basis (Verbaarschot et al., 2015). The averaging of the signal over many trials causes a loss of dynamic information, and the loss of key features unique to individual trials and individual participants. In both instances, the RP starts ∼ 200 ms before the action; which is relatively later when compared to the RP of the original Libet paradigm (Libet et al., 1983). The machine learning approach appears to detect activity earlier than the RP described in the literature, as well as being confirmed in this study; thereby making machine learning more informative about the decision / action initiation than the RP alone. The cluster based permutation test on the single channel data found a significant difference between −70 and −200 ms, a result consistent with the lateralised readiness potential. The cluster based analyses on the entire montage found clusters with significant differences between the timestamps −3250 and −4500. These significant differences confirm there are features specific to each decision within the EEG data leading up to the action, however the machine learning performance is superior to this. This shows that there is limited neural information present in the RPs, and as such cannot be used to contribute to the understanding of neural preparatory processes of motor activity.

It is for that reason the aim of this study was to accurately classify a decision (left or right), by employing a supervised learning deep neural network (i.e. a convolutional neural network) using only pre-conscious EEG data as the input. This was achieved using a convolutional neural network with a test classification accuracy of ∼ 99 % up to 4.45 s before the movement. This near perfect accuracy was achieved on an individual trial basis - an excellent indicator of the model’s performance. This accuracy drops to 55 % and 50 % when the windows extend from five to seven seconds before the action. These results are no better than chance. This suggests the decision arises in the frame between 5000 and 4450 ms. This difference cannot be entirely attributed to the difference in the number of training samples, as this difference is not as significant as the difference in accuracies for the respective windows Furthermore, the CNN model was able to predict each individual action made by the participant. This result supports the claim that the neural decisions are set before the time people report their conscious awareness of the decision. In addition, it is able to do this on an individual trial basis using data that is more representative of the brain’s function. This improved representation of brain activity is important, as it maintains the complex and dynamic nature of the brain. With each decision, our brains present a little differently, and these differences are preserved in the data inputted to the CNN model. The success of the CNN can be attributed to there being features relating to decision-making more than three seconds before conscious awareness. This is an improvement on the RP, as it only presented a few hundred *milliseconds* before the action and conscious awareness, which doesn’t account for significant pre-conscious involvement.

Although this study did not manage to predict as early as in the experiments of Soon *et al*. (2008, 2013), it is still able to predict with classification accuracies greater than that of Soon *et al*. (2008). Soon *et al*. (2008) achieved accuracies of 60 % up to seven seconds before conscious awareness which is close to chance. However, some participants in this study decided to move very quickly and as a result there were very few window periods of 7-10 seconds for each participant to input into the CNN model to ensure adequate training samples.

Another major criticism was addressed through the addition of a choice, as the RP has been argued to reflect neural activity of no more than an urge (Soon et al., 2008), or the product of unspecific neural activity (Trevena & Miller, 2010). In investigating free will, as it is used in the context of this study, there is the explicit need to have been able to do otherwise, given a rerun of the exact same scenario (Dias & Lavazza, 2016). This follows on from the work of Soon *et al*. (2008, 2013) in that the participants were given an explicit choice in each trial. This method is still limited to a degree, in that there is no consequence (a third requirement of a free choice) and this experiment doesn’t allow for replication of a real-world scenario such as that introduced in the work of. Further research into the addition of consequence, such as that of Maoz *et al*. (2019), is a promising perspective for future research.

The success of the CNN model gives conclusive evidence that there are features in the pre-conscious EEG data relating to a decision. These features are identifiable on an individual trial basis, unlike the readiness potential of Libet *et al*. (1983) and are specific to either decision (“left” or “right”). The features are present in the pre-conscious with the participants having had the option to choose an alternative path – which aligns with the definition of volitional actions used in this study (Dias & Lavazza, 2016). This further supports the notion that the agency for our decisions belongs to the pre-conscious. This supports the conclusions of Fried *et al*. (2011); Libet *et al*. (1983); Soon *et al*. (2008, 2013) and provides evidence to the notion that there is neural information highly predictive of a free choice present before the subjective report of conscious awareness of a decision to move by participants here demonstrated to be up to 4.5 seconds before the movement. This result has put machine learning’s foot solidly in the door of contributing to the empirical neuroscientific studies into volition.

The length of the frames of EEG data input into the model is a limitation of this study. Varying lengths of frames of EEG data from before the moment of conscious awareness were fed into the model to make the classification, resulting in imprecise timing in terms of exactly when the feature arose before conscious awareness of a decision to move. The core of the Libet paradigm is centred around the timing of events, making the consideration of temporal resolution of events important. Further limitations extend to the decision in this experiment was inconsequential, an important component of a volition decision (versus a reaction, for example) as well as the limitation in the reduction in the number of samples available for input as the length of the window increased. Fewer samples were available for training the model for the longer windows, as the time before an action carried out was variable and not controlled during the experiment.

Despite successfully using machine learning for classifying actions using only EEG data before conscious awareness of a decision to move, there is still little understanding of the mechanisms and features the model identified during its training. Further research, through the visualisation of the CNN model’s learned features, is necessary to understand the exact mechanisms governing the pre-conscious. Visualising the model’s learned features is a means of understanding what features the model used, where these features are in the brain, and the exact time they arose.

## 5. Conclusion

This study has shown machine learning can be used to develop some understanding the neural information relating to when free decisions arise in the brain. There is still much to be done in terms of improving the experiment itself, and the methods of analysing the data. However, this study was a proof of concept and aimed to establish whether machine learning can contribute to the current literature. The deep learning neural network, viz. the convolutional neural network employed, in this study achieved almost perfect accuracy in classifying an action using pre-conscious EEG data. This shows that machine learning, with a specific focus on deep learning is an important tool to include in all research in this domain going forward.

